# Exotic catenulid flatworms (Platyhelminthes, Catenulida) and where to find them in a temperate climate – a field study in a botanic garden

**DOI:** 10.64898/2026.08.06.743217

**Authors:** Katarzyna Tratkiewicz, Monika Sysiak, Marcin Zych, Ludwik Gąsiorowski

## Abstract

Catenulids are free-living flatworms, common in eutrophic freshwaters such as ponds, ditches, or peatbogs, with most of the diversity described from tropical regions to date. Although the majority of the species have been described from warmer climates, most molecular studies have been done on specimens from temperate zones in Europe. We addressed this gap by sampling for exotic species in localities available in a temperate climate. In this study, we investigated catenulid diversity in the greenhouses at the University of Warsaw Botanic Garden and recorded two species known only from tropical areas (*Stenostomum paraguayense* and *Suomina evelinae*) and one exotic species recorded previously from a greenhouse in Poland (*Stenostomum corderoi*). Additionally, in the latter species, we provide evidence for environmentally induced coloration of sensory pits, which has not been reported thus far. We placed the collected species on a phylogeny using barcoding of 18S, 28S, and COI genes and retrieved paraphyly of the family Catenulidae, with *S. evelinae* forming a sister group to the genus *Paracatenula*, and hence we propose a revision of its systematic position. In total, we recorded six species, including three with a wide cosmopolitan distribution *(C. turgida*, *S. grande* and *S. tuberculosum*), and provided sequences for five of them, three of which had no previous molecular records *(S. paraguayense, S. evelinae* and *S. corderoi)*. Thus, we confirm that greenhouses represent an important source of exotic species for taxonomic work on microscopic invertebrates.

## Introduction

Catenulida is an order of free-living microscopic flatworms, occupying a sister position to all of the other platyhelmints (Larsson and Jondelius 2008, Egger et al. 2015, Laumer et al. 2015). Although some families are predominantly marine, most catenulids live in eutrophic freshwaters (Nuttycombe and Waters 1938, Marcus 1945, Larsson and Willems 2010). Most of their diversity to date was reported from tropical habitats, with over half of the species of the largest family, Stenostomidae, noted from South America (Marcus 1945, van der Land 1970, Noreña Janssen 1995, Noreña et al. 2005, Damborenea et al. 2011). Catenulids are found in a diverse range of habitats from lakes and peatbogs (Nuttycombe and Waters 1938, Larsson and Willems 2010) to more specialized environments such as rice fields (Yamazaki et al. 2012) and inside the vase-shaped water-accumulating leaf rosettes in some bromeliads (plant family *Bromeliaceae*) (Marcus 1945).

Despite their cosmopolitan occurrence in various environments, catenulids remain taxonomically understudied because of difficulties with their identification. In general, they lack hard reproductive structures, used extensively in the taxonomy of other flatworms (Brusa et al. 2020). Additionally, they exhibit phenotypic plasticity (Nuttycombe 1956, Rosa et al. 2015, Tratkiewicz et al. 2026), making characters such as shape or size unreliable. Furthermore, particular authors give unequal weight to diagnostic traits, making comparison of species from different geographic regions difficult. All of the above reasons result in a scarcity of comprehensive taxonomical works on catenulids (Nuttycombe and Waters 1938, Marcus 1945, Luther 1960, Kolasa and Young 1974, Noreña et al. 2005), with most of the molecular studies to date performed in temperate climate zones in Europe (Larsson and Jondelius 2008, Diez and Schmidt-Rhaesa 2024, Tratkiewicz et al. 2026). This causes a gap in our understanding of catenulid diversity, which we tried to address in this study by sampling for tropical species at easily available localities, i.e., greenhouses at the botanic garden.

Botanic gardens form a very specific environment, often described as ‘tropical islands’ in a temperate climate, where many species are introduced under uncontrolled conditions via components such as soil or plants (Zawierucha et al. 2013, Kolicka et al. 2015). Among the organisms that can thrive in such settings are catenulids, which reproduce asexually through a process called paratomy (Moraczewski 1977, Palmberg 1990, da Rosa and Loreto 2024, Gąsiorowski et al. 2025), allowing even a single individual to establish a new population. In fact, catenulids have been previously researched in greenhouse conditions by Kolasa (1973), who reported several species of *Stenostomum*, *Catenula* and others from a botanic garden in Poznań, Poland (Tab. 1). One particularly well-studied microhabitat found in greenhouses is the water-filled vase-shaped leaf rosettes of *Bromeliaceae* plants, which have attracted scientific attention for over a hundred years (eg. Scott 1912, Richardson 1999, Lopez et al. 2009, Lounibos and Frank 2009, Medeiros et al. 2024). Although some catenulid species have been recorded from bromeliads in natural environments (Marcus 1945), the most recent study of bromeliad fauna in greenhouse conditions by Kolicka et al. (2016) found several testate amoebae, gastrotrichs, rotifers and other invertebrates in these microreservoirs, but no flatworms were identified, a gap we also aimed to address.

**Table 1.**
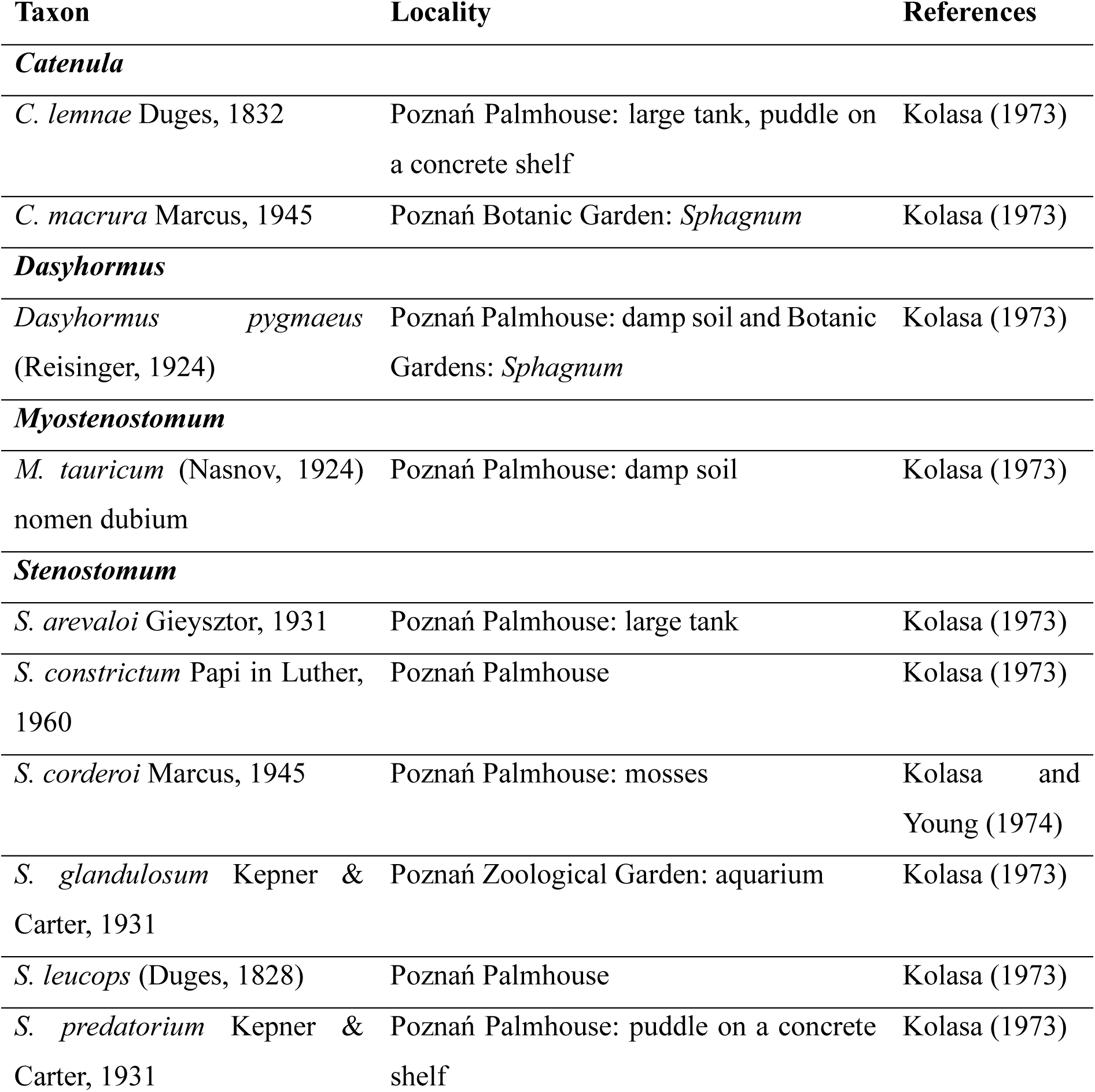
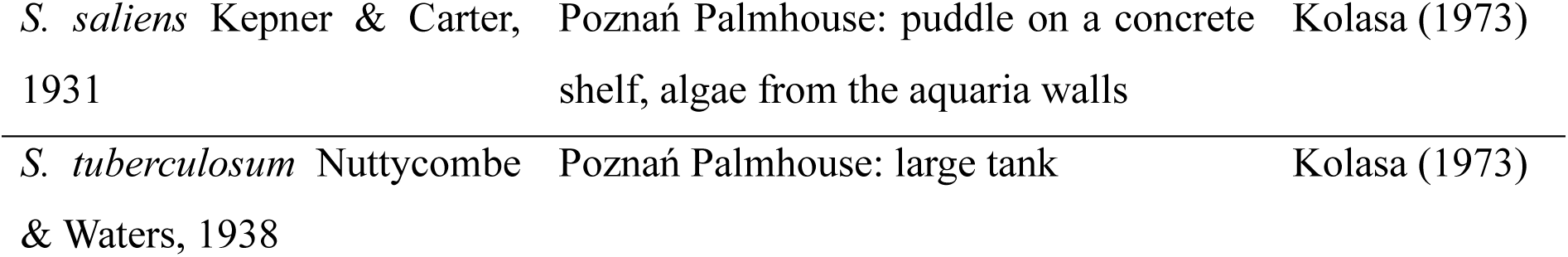
Revision of Catenulida representatives previously reported from botanical gardens.

In this study, we investigated the presence and distribution of catenulid flatworms at the University of Warsaw Botanic Garden greenhouses, studying catenulid faunas both in artificial water bodies and associated with the bromeliad plants. We provided sequences of three markers (18S, 28S and COI) for five species, three of which lacked any prior molecular data. We reconstructed their phylogeny and placed them on the Catenulida tree, showing that greenhouses can serve as an important source of microinvertebrate species for taxonomic studies.

## Methods

### Study site

Sampling was conducted at the University of Warsaw Botanic Garden (Warszawa, Poland). Founded in 1818, it is the third oldest botanic garden in Poland, however, its greenhouses were reopened in the late 1940s, after being destroyed in 1944. In this study, samples were collected from three different buildings – Subtropical, Palm, and the backside of the Tropical greenhouse (Fig. 1).

**Figure 1.**
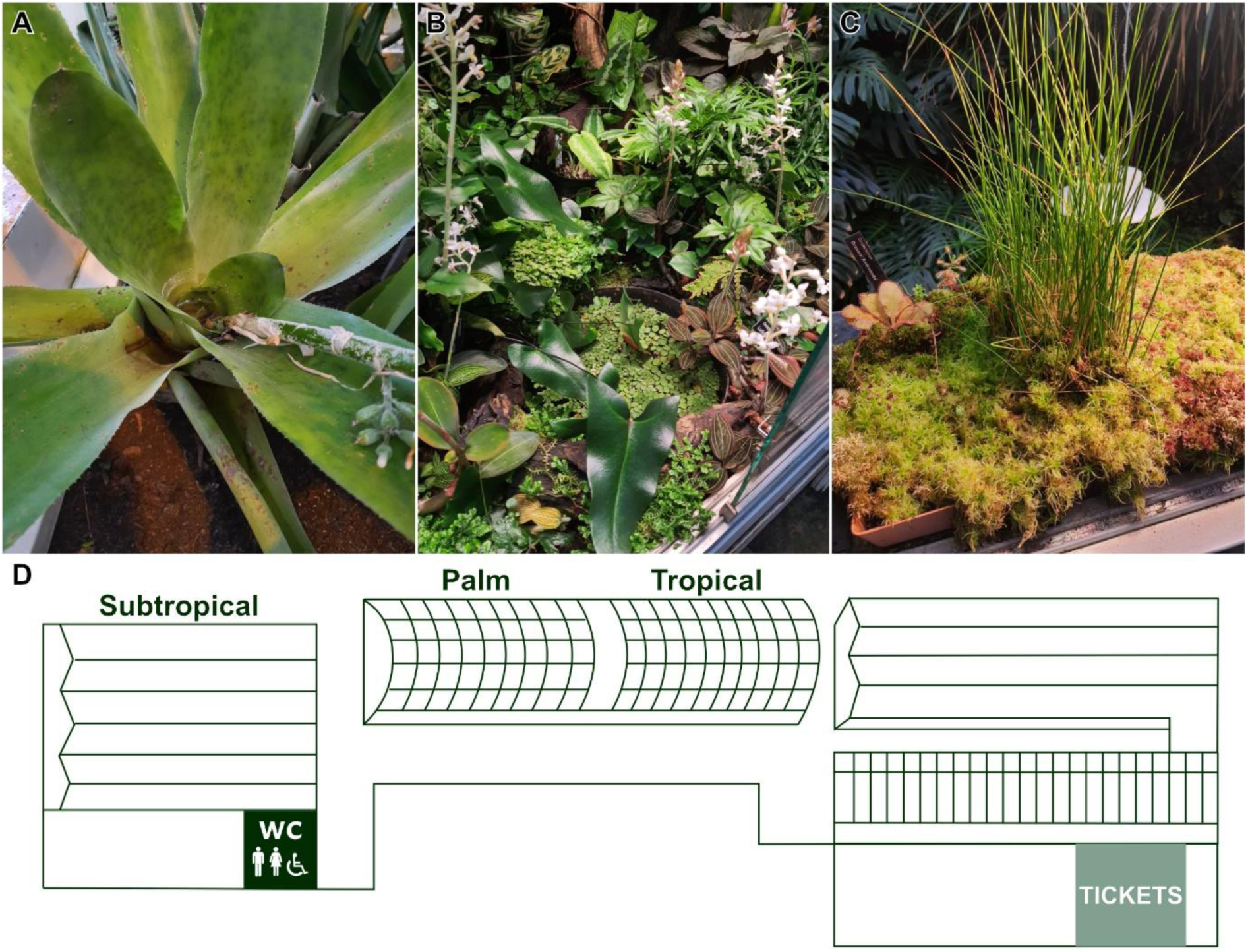
Examples of studied reservoirs and a map of sampling sites. **A.** Inside of *Aechmea lueddemanniana* (*Bromeliaceae*); **B.** Tank in a cabinet; **C.** *Sphagnum* sp.; **D.** Map of sampling sites.

### Animal collection and documentation

Sampling was conducted by hand. The water samples were taken into 50 mL Falcon tubes. In the case of *Bromeliaceae* reservoirs, the water was collected using a plastic Pasteur pipette with a wide tip. For *Sphagnum* extractions, fragments of plants were collected into Ziploc bags, then submerged into distilled water and extracted using the squeeze method on a 70μm sieve. During the sampling, an oxygen probe (YSI ProODO YSI®) was used to measure the temperature and the oxygen concentration of samples (if possible). Afterward, the pH was checked with a pH meter (CP-505 ELMETRON®). Details on environmental parameters and species identified in each sampling site are provided in Table 2.

**Table 2.**
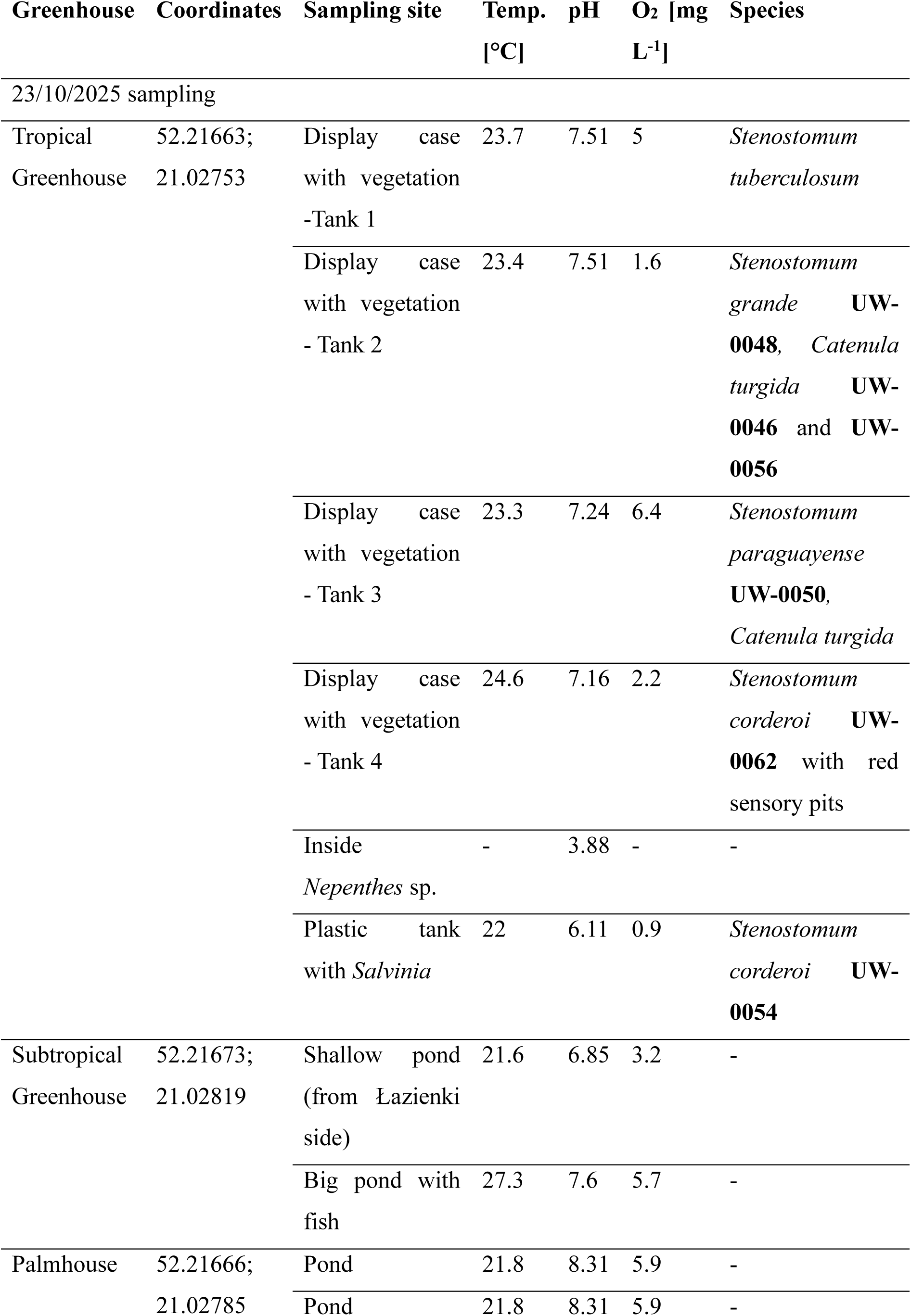
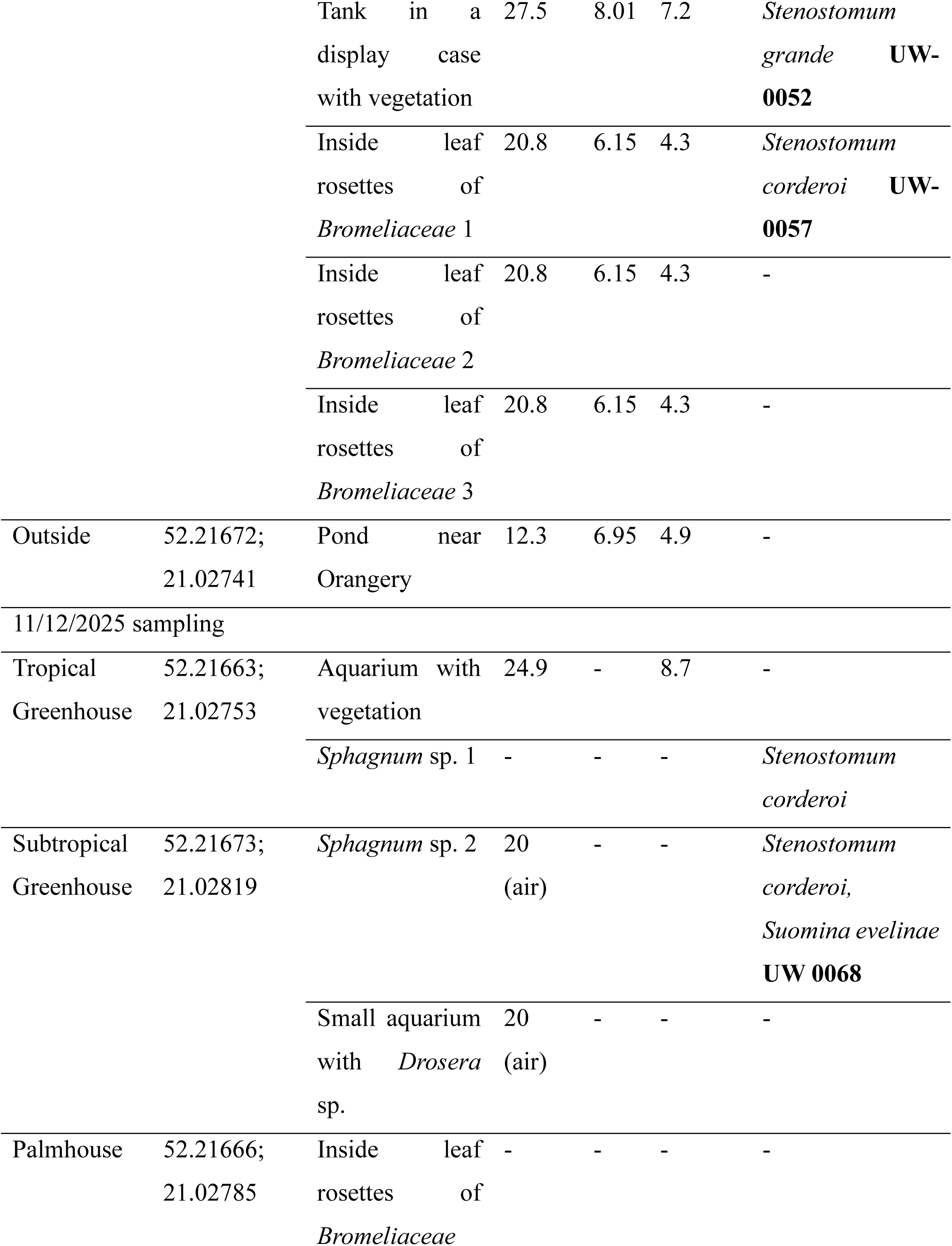
Catenulid taxa identified in different sampling sites.

Samples were examined in plastic Petri dishes under the stereomicroscope. Animals were sorted with automatic pipettes and imaged with a Nikon Eclipse NI-SSR microscope equipped with Nomarski contrast optics and a Nikon DS-Ri2 camera. Contrast and brightness of the photographs of the animals were adjusted in the image analysis software Fiji (Schindelin et al. 2012).

### DNA extraction, PCR amplification and sequencing

After photographic documentation, the DNA was extracted from single worms using Chelex 100 resin (Bio-Rad, Hercules, California, USA) – following the procedure from Tratkiewicz et al. (2026). Then, the PCR was performed in a mixture consisting of 2.5 μl of isolated DNA, 0.25 μM of forward and reverse primers, and 12.5 μl of NZT Taq II 23 Green Master Mix (NzyTech, Lisboa, Portugal), in a final volume of 25 μl. The genes were amplified with the following primers: 4fb + 1806R for 18S gene, LSU5 + L1642R for 28S, and COI5B + COI3B for COI (Larsson and Jondelius 2008). The PCR protocol began with 3 minutes of denaturation at 95°C, followed by 35 cycles comprising 30 s at 94°C; 30 s at 50°C (18S), 53°C (28S) or 43°C (COI); and 30 s at 72°C, followed by a final extension at 72°C for 5 min. The PCR products were purified with Syngen PCR Mini Kit (Syngen Biotech, Wrocław, Poland). Purified DNA was sequenced using the Sanger method by Genomed S.A. (Warsaw, Poland). For the COI gene, the COI5B primer was used for sequencing; the 18S and 28S genes were sequenced with both forward and reverse primers. Sequences were assembled and trimmed using Geneious v2026.0.2. Then the sequences were deposited in GenBank under accession numbers PZ143194-PZ143204; PZ164137-PZ164143 and PZ210113-PZ210119.

### Phylogenetic analyses

A dataset combining reference sequences with those obtained in the current study was assembled (Tab. S1). The sequences were aligned with MAFFT v7.490 (Katoh and Standley 2013) with the automatic (Auto) algorithm setting. Ambiguously aligned regions were trimmed with the Mask Alignment tool in Geneious using a 50% gaps setting. The final concatenated alignment of 3580 nucleotides was partitioned into 18S, 28S, and COI partitions. The best substitution model for each marker was estimated with ModelTest-NG v0.1.7 implemented in RAxML GUI v2.0.16 (Edler et al. 2021), and then the Bayesian phylogeny was reconstructed using MrBayes 3.2.7 (Ronquist et al. 2012) employing default priors and the same partitioning scheme. Two simultaneous analyses were conducted, each comprising 2 million generations with one cold and three heated chains, sampling every 1000 generations. Tracer v.1.5 (Rambaut et al. 2018) was utilized to assess the convergence of the Markov Chain Monte Carlo (MCMC) chains.

## Results

### Morphology of sampled species

Our collection yielded nine barcoded specimens, representing five morphotypes, belonging to three genera: one species of *Catenula*, one species of *Suomina,* and three species of *Stenostomum*.

### *Catenula turgida* (Zacharias, 1902)

#### *Sampling sites:* Display case with vegetation - Tank 2 and 3

A *Catenula* reaching 400 μm in length. The body shape is conical, with a narrowing posterior part (Fig. 2A). The head is clearly separated from the rest of the body, with a slight tripartition (Fig. 2A). In our specimens, we could not observe a statocyst. At the base of the head, a ring of cilia is present (Fig. 2D) with four furrows on the ventral side. The ciliated ring is followed by a small triangular mouth opening and a long pharynx with cilia arranged like a comb (Fig. 2B). The gut does not reach the posterior end of the body, which is blunt, without a tail. The protonephridial canal is visible throughout the whole length of an animal and ends in the caudal region. In the epidermis, around 3 μm long rod-shaped rhabdites occur, grouped into sets of a few (Fig. 2C). The animal is covered by a uniform layer of cilia, with longer, stiffer ones distributed sparsely but evenly around the body (Fig. 2B).

**Figure 2.**
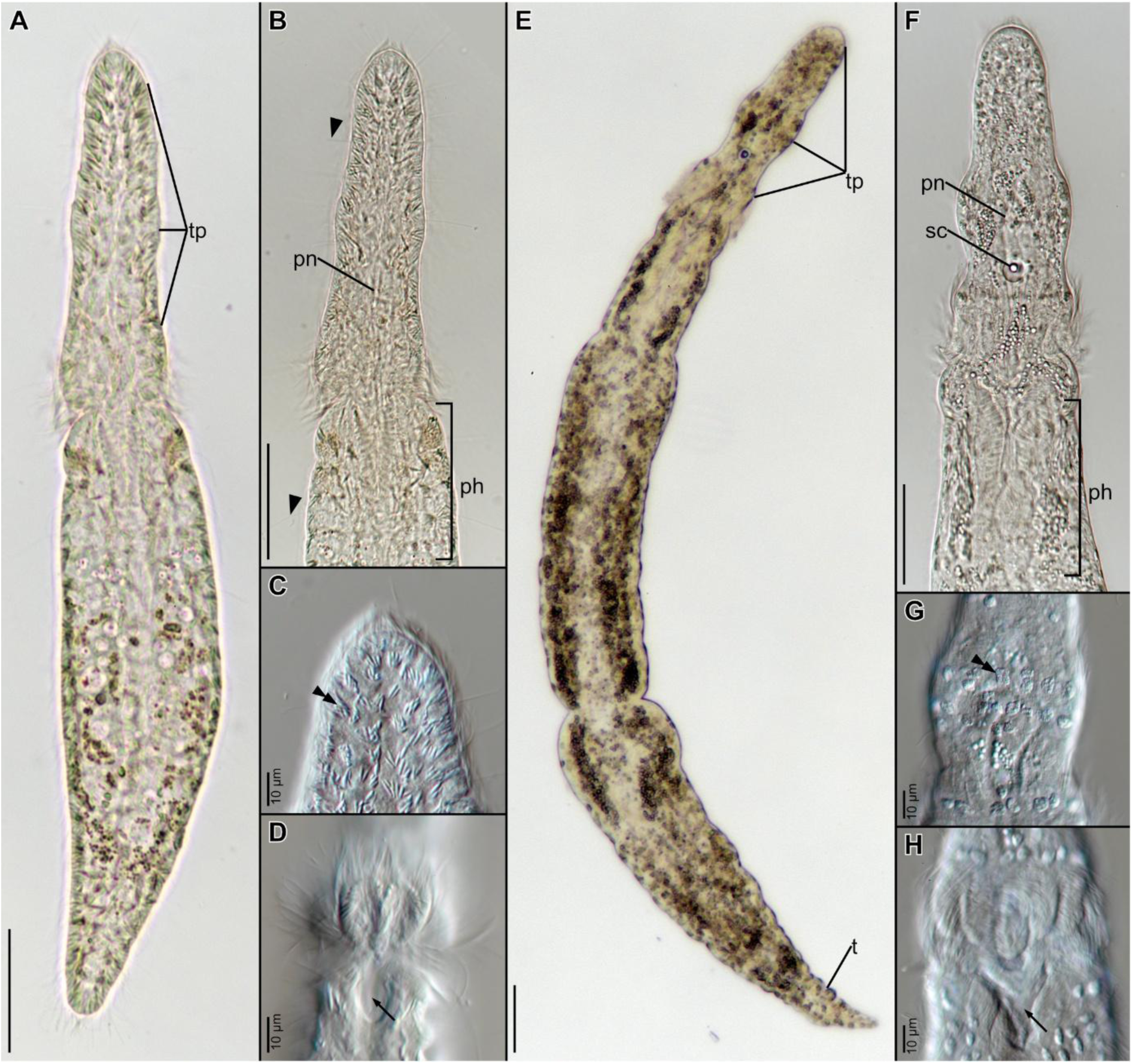
Morphology of a *Catenula turgida* (**A-D**) and *Suomina evelinae* (**E-H**). **A.** Overview of the animal; **B.** Overview of the head; **C.** Magnification of the rhabdites in the epidermis; **D.** Magnification of the ciliated ring in the mouth region; **E.** Overview of the animal; **F.** Overview of the head; **G.** Magnification of the epidermal inclusions; **H.** Magnification of the ciliated ring in the mouth region. Abbreviations: ph, pharynx; pn, protonephridium; sc, statocyst; t, tail; tp, tripartition of the head; arrow, mouth opening; arrowhead, long stiff cilia; double arrowhead, rhabdites/epidermal inclusions. Scale bars: 50 μm (**A, B, E, F**); 10 μm (**C, D, G, H**).

The worms can be cultured with some difficulty on Chalkley’s Medium with wheat grain at 20°C. In the cultures, they swim freely in the water column or crawl in the sediment at the bottom of the dish and prey on unidentified bacteria.

*Distribution:* Germany, *Sphagnum* in moorland near Plön (Zacharias 1902); Brazil, bromeliads and stream in São Paulo (Marcus 1945); Finland, raised bog in Tvärminne (Luther 1960); Sweden, *Sphagnum* from various locations (Larsson and Willems 2010); Brazil, wetland in Rio Grande do Sul (Braccini et al. 2017) – however, the general body shape of the latter is more reminiscent of *C. sawayai*.

### Suomina evelinae Marcus, 1945

#### Sampling site: Sphagnum sp. 2

Animals reach a length of 800 μm in the two-zooid stage; chains longer than two zooids were not observed. The body is cylindrical, with a narrowing and pointy posterior end (Fig. 2E). The head is not separated from the body; however, it has a clearly distinguishable tripartition (Fig. 2E). The first thickening at the head appears at the anterior part, followed by another in the middle, with the last one before the mouth opening. Between the second and third thickening, there is a statocyst with a single, spherical, greenish statolith (Fig. 2F). On each of the three parts, rounded granular epidermal inclusions are present (Fig. 2G). The third thickening is surrounded by a ciliated ring (Fig. 2H) split into four furrows on the ventral side. The mouth opening is located behind the ciliated ring, followed by a pharynx shaped like a comb, similar to the one in *C. turgida*. The gut was brownish in our specimens, but the inner lining of the intestine was not clearly distinguishable. The posterior end of the body is creased and resembles a tail (Fig. 2E). The cilia layer is uniform and covers the whole animal; no stiff cilia were observed.

The worms are very easily culturable on Chalkley’s Medium with wheat grain at 20°C. In the cultures, they swim freely in the water column or crawl in the sediment at the bottom of the dish and prey on bacteria and small protozoans.

*Distribution:* Brazil, bromeliads from São Paulo, Pinheiros River meadow (Marcus 1945); Brazil, wetland in Rio Grande do Sul (Braccini et al. 2017).

### Stenostomum corderoi Marcus, 1945

#### *Sampling sites:* Display case with vegetation - Tank 4 (red pits), Plastic tank with *Salvinia* sp., Inside leaf rosettes of Bromeliaceae 1, *Sphagnum* sp. 1, *Sphagnum* sp. 2

Single worms reach up to 600 μm in length (Fig. 3A), often occurring in longer chains consisting of up to five zooids. The body is cylindrical, narrowed in the anterior part and rounded at the posterior end with a noticeably wider pharyngeal region. The sensory pits are small and round, enclosed in the head, located in front of the brain lobes (Fig. 3B). The animals lack refracting bodies. The pits and brain arrangement are similar to those of *S. brevipharyngium* and *S. saliens,* but the species is distinguishable by a pharyngeal region. The mouth opening is V-shaped, extending to half of the pharynx, with the rest of the pharynx having a characteristic creased surface (Fig. 3B). Excretophores appear in the gut, which has a characteristic wavy lining (Fig. 3C). The ciliary coat is uniform, with a few stiffer cilia at the anterior and posterior parts of the worm. At the very posterior part, an additional cluster of normal cilia appears (Fig. 3C).

**Figure 3.**
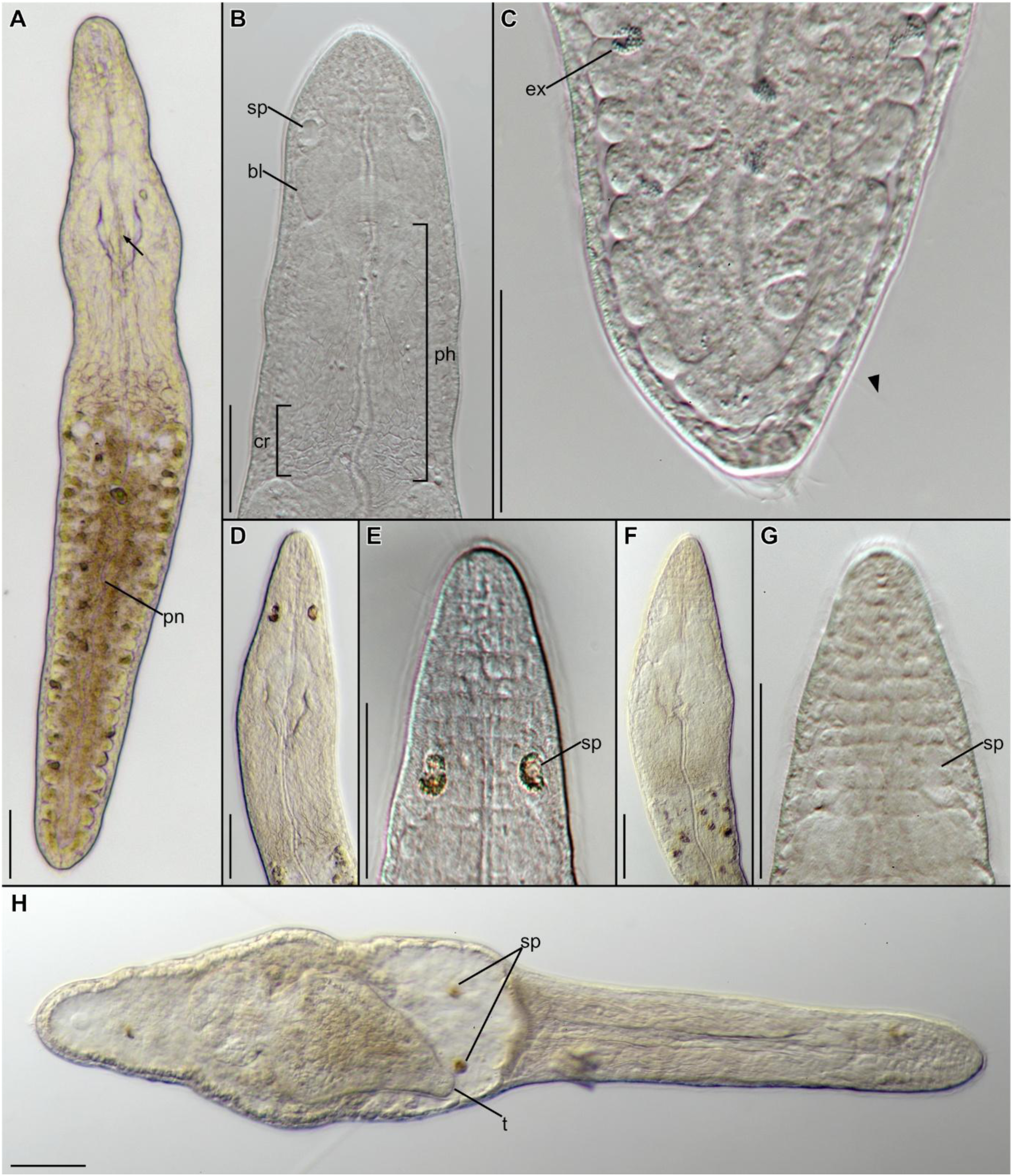
Morphology of a *Stenostomum corderoi* from Tank with *Salvinia* (**A-C**) and the individuals with red sensory pits from Cabinet - Tank 4 (**D-H**). **A.** Overview of an entire worm; **B.** Magnified view of the head; **C.** Magnified view of the caudal region; **D.** Overview of the head of the individual with red sensory pits from Cabinet - Tank 4; **E.** Magnification of the sensory pits of the same individual; **F.** Overview of the head of the morphotype from Cabinet - Tank 4 put in water from Tank with *Salvinia* for 24h; **G.** Magnification of the sensory pits of the same individual; **H.** Cannibalism in *S. corderoi*. Sensory pits and tail of the predated conspecific worm are visible inside the gut. Abbreviations: bl, brain lobe; cr, creased area of the pharynx; sp, sensory pit; ex, excretophores; ph, pharynx; pn, protonephridium; arrow, mouth opening; arrowhead, long stiff cilia. All scale bars equal 50 μm.

Some individuals found in Cabinet - Tank 4 exhibited red sensory pits, noticeable even under the dissecting microscope (Fig. 3D-E). Initially thought to represent a different species, these individuals were, upon closer examination, found to exhibit morphology consistent with *S. corderoi*. Animals were separated and moved into filtered water from another location: a tank with *Salvinia*. After 24 h in the water from the *Salvinia* tank, the coloring of the pits disappeared (Fig. 3F-G). Later, another set of experiments was conducted. Two animals from each location – both Tank 4 and the *Salvinia* tank were separated and put in the 4-well plate. One of the animals was kept in the original filtered water, and the other one was put in the filtered water from the other location. After one day in changed water, the animals with red pits (Tank 4) lost the coloration, and the animals that exhibited normal pits (Tank with *Salvinia*) gained the red color. The color of the pits in control animals in filtered water from the same location did not change.

The worms are easily culturable on Chalkley’s Medium with wheat grain at 20°C. They swim freely in the water column and prey on small protozoans. When starved, a few instances of cannibalism were observed (Fig. 3H).

*Distribution:* Brazil, bromeliads from São Paulo (Marcus 1945); France (De Beauchamp 1948); Poland, mosses from Poznań Palmhouse (Kolasa and Young 1974).

### *Stenostomum tuberculosum* Nuttycombe & Waters, 1938

#### *Sampling site:* Display case with vegetation - Tank 1

A *Stenostomum*, reaching a length of 600 μm in two-zooid individuals (Fig. 4A). The head is triangular with a characteristic tubercle at the very top (Fig. 4B). Sensory pits are directed anteriorly, deeply enclosed in the head. The pits are connected to the posterior brain lobes, which are oval (Fig. 4B). The animals lack refracting bodies. The mouth opening is V-shaped and strongly muscular (Fig. 4A). The pharynx is relatively short with fine granular gland cells (Fig. 4B). The anterior end of the gut does not extend to the caudal end, which is narrower than the rest of the body. The protonephridium is straight, with a terminal opening at the very end of the animal.

**Figure 4.**
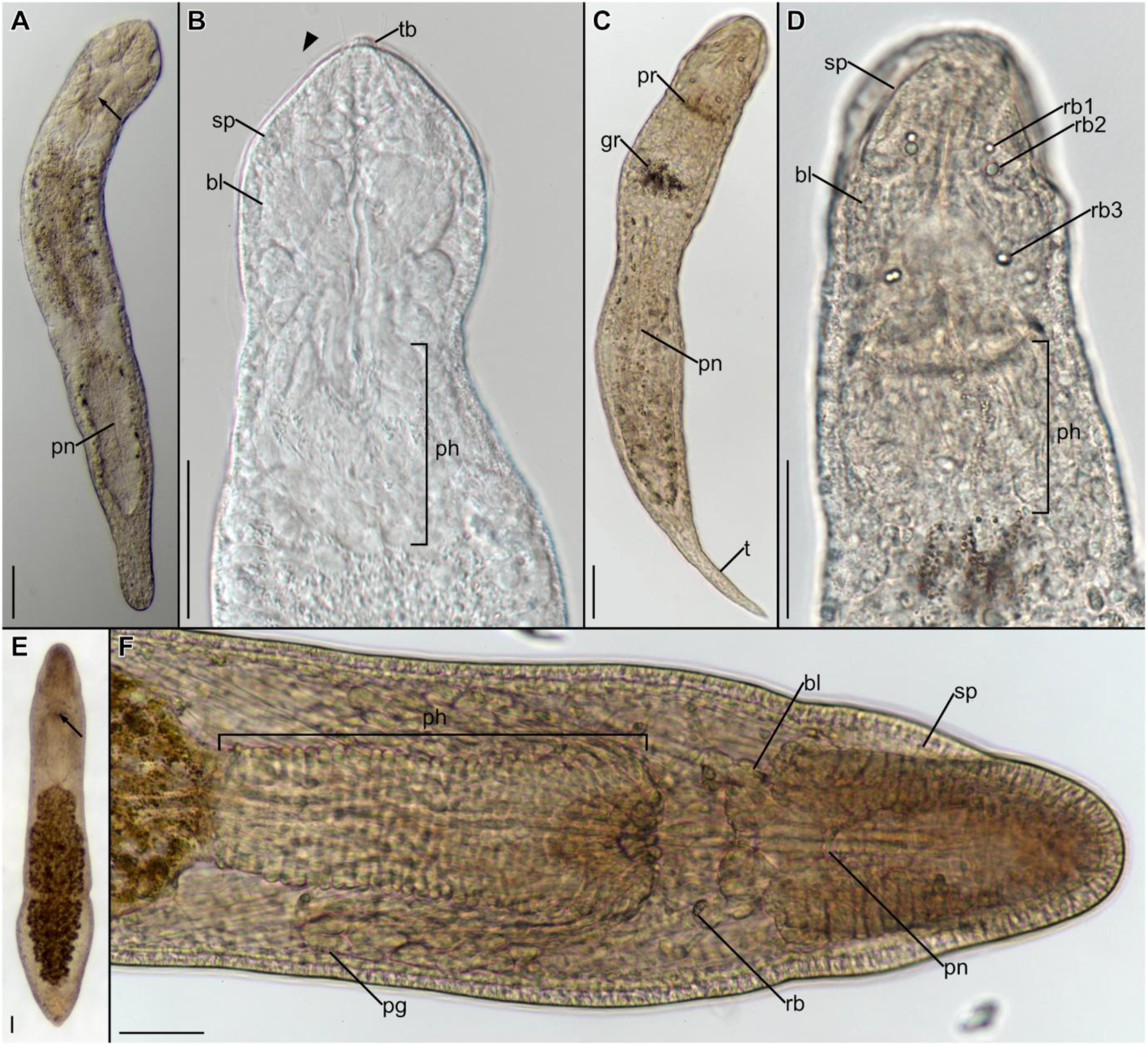
Morphology of a *Stenostomum tuberculosum* (**A-B**), *S. paraguayense* (**C-D**) and *S. grande* (**E-F**). **A.** Overview of the *S. tuberculosum*; **B.** Magnification of the head of the *S. tuberculosum*; **C.** Overview of the *S. paraguayense*; **D.** Magnification of the head of the *S. paraguayense*; **E.** Overview of the *S. grande*; **F.** Magnification of the head of the *S. grande*. Abbreviations: bl, brain lobe; sp, sensory pit; gr, gut ring; pg, pharyngeal gland; ph, pharynx; pn, protonephridium; pr, pharyngeal ring; rb, refracting body; t, tail; tb, tuberculum; arrow, mouth opening; arrowhead, long stiff cilia. All scale bars equal 50 μm.

The worms survived for only a few days in Chalkley’s Medium with wheat grain before the culture collapsed. We did not succeed in further attempts of culturing them.

*Distribution:* US, Mountain Lake, Virginia and pond at Athens, Georgia (Nuttycombe and Waters 1938); Brazil, streams in São Paulo (Marcus 1945); Poland, tanks in Poznań Palmhouse (Kolasa 1973); Surinam, Suriname River near Paramaribo (van der Land 1970); Germany and Finland (Lanfranchi and Papi 1978); Argentina, Buenos Aires (Noreña et al. 2005); Japan, Ikeura rice field (Yamazaki et al. 2012); Brazil, ESEC Taim water channel (Reyes et al. 2021).

### Stenostomum paraguayense (Martin, 1938)

#### *Sampling site:* Display case with vegetation - Tank 3

A *Stenostomum* reaching the length of 500 μm in a single worm. The body is cylindrical with a distinguishable tail and a rounded anterior part (Fig. 4C). The sensory pits are elongated and connected to the anterior, metameric brain lobes (Fig. 4D). There are three pairs of refracting bodies. The first pair occurs at the middle of the length of the pits, the second at the end of the pits, and both consist of a single sphere. The third pair is connected to the end of posterior brain lobes and is made from two connected spheres (Fig. 4D). The mouth opening is small and slender, not easily recognizable on the photos; however, it is accompanied by a very distinguishable ring of darker cells (Fig. 4C). The short pharynx passes into another ring of darker cells at the beginning of the intestine. The presence of these two dark rings is characteristic of the species. The body is yellowish with a uniform layer of cilia. The gut does not reach the caudal end of the body, which ends in a long tail (Fig. 4C).

This species is highly similar to *S. arevaloi;* however, according to Marcus (1945), it differs in width, mouth shape, gland cells at the beginning of the intestine, and dorsocaudal appendix. We did not observe any easily distinguishable appendix, but according to Marcus, the structure is not always present.

The animal can survive for a few days in Chalkley’s Medium, fed on rotifers *Lecane inermis*, but the worms progressively decreased in size and died, likely due to starvation. We did not succeed in attempts of culturing them.

*Distribution:* Brazil, São Paulo tanks and aquariums, Pinheiros River meadow, Canindé lake (Marcus 1945); Argentina, Córdoba (Adami and Damborenea 2021) – described as *S. arevaloi* but very likely *S. paraguayense*.

### Stenostomum grande Child, 1902

#### *Sampling sites:* Display case with vegetation - Tank 2, Display case with vegetation in a Palmhouse

A large *Stenostomum* reaching the length of 1100 μm in a two-zooid individual. The body shape is cylindrical and compact with pointy anterior and posterior parts (Fig. 4E). The sensory pits are elongated and open to the environment, similar to the ones in *S. paraguayense*. They are enclosed shallowly in the anterior brain, which is composed of many metamers (Fig. 4F). The posterior brain is made from four lobes with the posterior ones connecting through protrusions with refracting bodies, which are tear-shaped and consist of many granules (Fig. 4F). The mouth opening is small but clearly visible and followed by a long pharynx, with strong circular musculature and a large number of glandular cells along the anterior two-thirds of its length (Fig. 4F). The gut is dark brown and does not reach the caudal end of the body, which ends in a small, triangular tail (Fig. 4E). The protonephridium is clearly visible, even at the dissecting scope, and ends after the gut, but does not reach the posterior extremity of the body.

The worms can be cultured on Chalkley’s Medium with wheat at 20°C, where the animals stay at the bottom of a dish, feeding on decomposing organic matter.

*Distribution:* US, New York and Massachusetts (Graff 1913); Russia, Kola Peninsula (Nassonov 1924); US, Virginia (Nuttycombe and Waters 1938); Brazil, São Paulo (Marcus 1945); Surinam, Onverwacht (van der Land 1970); Poland, heated Konin Lakes (Kolasa 1977); US, New York (Kolasa et al. 1987); Japan, Ikeura rice field (Yamazaki et al. 2012); Argentina, Buenos Aires (Noreña et al. 2005); Lake Mangueira and Lake Nicola (Reyes et al. 2021).

### Phylogenetic position of sampled species

Bayesian analysis recovered a monophyletic Catenulida, divided into two clades (Fig. 5). The first clade consisted of the family Stenostomidae (*Stenostomum* and *Rhynchoscolex*), and the second of Catenulidae, Paracatenulidae and Retronectidae. The family Catenulidae was recovered as paraphyletic, as specimens identified morphologically as *S. evelinae* (Catenulidae) grouped with high support with *Paracatenula* (Paracatenulidae), and not with the members of the genus *Catenula* (Catenulidae). Specimens of *C. turgida* grouped with the conspecifics worms from Sweden. *Stenostomum* clade shows a topology congruent with that of Tratkiewicz et al. (2026), with four well-supported clades (Clades 1-4) and poorly resolved deep nodes. *Stenostomum paraguayense* grouped closely with *S. arevaloi* Gieysztor, 1931 and *S. glandulosum* Kepner & Carter, 1931, forming a sister group to all the remaining *Stenostomum* (here referred to as Stenostomidae Clade 5). *S. corderoi* was placed within Stenostomidae Clade 4 (with *S. brevipharyngium* Kepner & Carter, 1931 as its closest relative) and *S. grande* in the Stenostomidae Clade 2 (grouping with *S. grande* from Japan).

**Figure 5.**
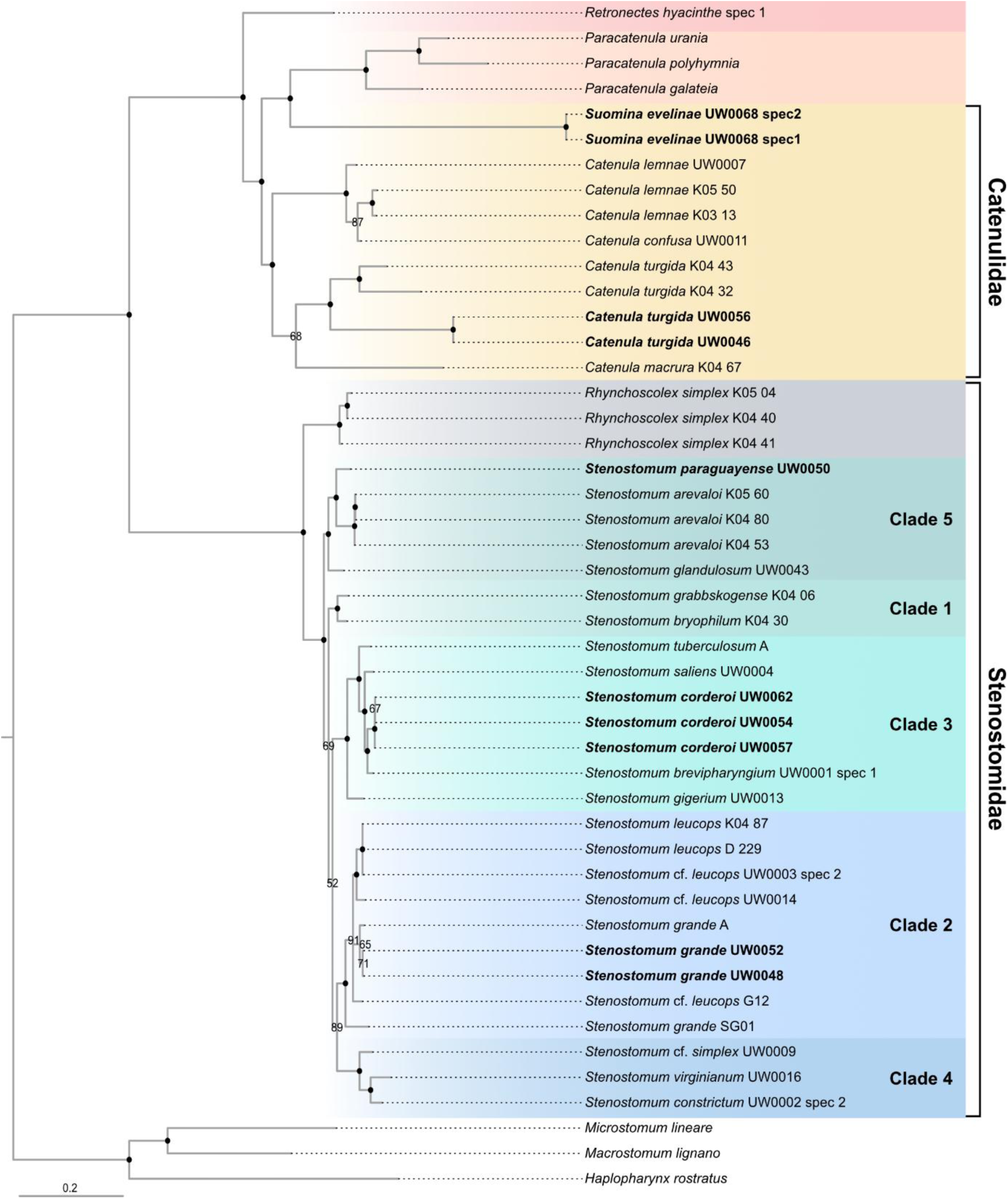
Consensus tree of Catenulida inferred with a Bayesian approach from the concatenated 18S, 28S and COI datasets. Filled dots indicate posterior probability > 95%, lower values are provided. Terminals in bold indicate newly sequenced individuals. Different genera are colored: *Retronectes* in red, *Paracatenula* in orange, *Catenula* in yellow, *Rhynchoscolex* in grey and *Stenostomum* in greens or blues. Different clades of *Stenostomum* following Tratkiewicz et al. (2026) and a newly defined Clade 5 are marked.

## Discussion

### Catenulid diversity in greenhouses

As reported in this study, greenhouses create suitable conditions for a variety of catenulid species. Although the routes by which they colonize botanic gardens remain unknown, tropical species may be transported with wet soil or plants. Borkott (1970) observed egg-laying in catenulids, but it is not known whether these eggs serve as dormant stages capable of withstanding extreme or unfavorable conditions, for example during long-distance transport. In total, we recorded six different catenulid species from the University of Warsaw Botanic Garden: three widely distributed around the world (*C. turgida*, *S. tuberculosum*, *S. grande*), one exotic species recorded previously from a greenhouse in Poland by Kolasa and Young (1974) (*S. corderoi*), and two known so far exclusively from tropical areas (*S. evelinae*, *S. paraguayense*). For the latter three, we provided the first reference molecular sequences for several barcodes. We report a similar species composition to that previously found in studies of botanic gardens. Besides *S. corderoi*, we recorded *S. tuberculosum*, which has been previously found in a Poznań Palmhouse, Poland (Kolasa 1973). Although we did not succeed in sequencing this species, its characteristic morphological feature (the tubercle) allowed us to identify it with a high degree of confidence. Another species recorded by Kolasa was *Dasyhormus pygmaeus*, which we argue is similar to *S. evelinae*; this is discussed further in the final part of the Discussion.

We did not observe any clear pattern regarding the environmental preferences of catenulids. They were found across a diverse range of pH from 6.11 to 7.51 and oxygen levels from 0.9 to 7.2 mg/l. No catenulids were found in the sample with the lowest pH recorded (3.88 in *Nepenthes*), suggesting they are not adapted to extremely low pH. *Stenostomum corderoi* was found in the tank with the lowest oxygen concentration in the study (0.9 mg/l), suggesting this species may be tolerant of near-anoxic conditions.

### Catenulids and bromeliads

The association between bromeliads and invertebrates has a long evolutionary history, estimated to have formed around 16 million years ago (Balke et al. 2008). Consequently, both the invertebrate and vertebrate fauna of bromeliads have been the subject of numerous studies (eg. Scott 1912, Richardson 1999, Lopez et al. 2009, Lounibos and Frank 2009, Medeiros et al. 2024), including a study of artificial bromeliad environments conducted in Poland by Kolicka et al. (2016), where turbellarian representatives were found but not identified to species or genus level. Some animal species are known to inhabit bromeliads exclusively, for instance the acari *Steneotarsonemus ananas* (Tryon, 1898) sensu Beer, 1954 or the gastrotrich *Chaetonotus* (*Hystricochaetonotus*) *furcatus* Kisielewski, 1991, but many species found in bromeliads are just opportunistic visitors. Although no catenulid species is known exclusively from bromeliads, we included these habitats in our study because some species have previously been recorded from them in the wild. Notably, Marcus (1945) reported in his revision of Brazilian microturbellarians, four catenulid species from bromeliads: *Suomina evelinae*, *Stenostomum anatirostrum*, *Stenostomum virginianum*, and *Stenostomum corderoi*. While the first three were also found in other habitats, the latter was reported exclusively from bromeliads. In our study, *S. corderoi* was likewise found in *Bromelia*, but it also occurred in small tanks and *Sphagnum* samples, making it the most common species across all sampling sites (Tab. 2). Although bromeliads were the only habitat reported in the original description of this species, it was later also recorded from mosses in a botanic garden (Kolasa and Young 1974). Given that bromeliads remain the only known natural habitat for *S. corderoi*, we speculate that the species is closely associated with them and may have been transported to greenhouses via *Bromeliaceae* plants, later colonizing other available habitats. Definitive conclusions, however, would require further sampling in its native range.

### Phenotypic plasticity in catenulids

Catenulids exhibit exceptional phenotypic plasticity, making traits such as body size or shape unreliable for taxonomy (Nuttycombe 1956, Rosa et al. 2015, Tratkiewicz et al. 2026). During this study, we found a small population of *S. corderoi* with red sensory pits in one of the tanks. Aside from the brightly pigmented pits, these animals were morphologically indistinguishable from specimens of *S. corderoi* found in the same cabinet but in different tanks. Subsequent sequencing confirmed they belonged to the same species, a finding further supported by the loss of pit coloration after the animals were transferred to water from a different tank. The pits are filled with a mucus-like substance (Ott 1892, Kepner and Cash 1915, Reuter et al. 1993, Gąsiorowski et al. 2023), and we therefore hypothesize that they can accumulate particulate matter or pigmented compounds dissolved or suspended in the water, causing the observed change in coloration. The genus *Xenostenostomum*, described by Reisinger (1976), was characterized by red, elongated pits, which could also have resulted from pigment accumulation, although this remains speculative as no photographs or drawings of this taxon exist. In light of our findings, we propose that pit coloration should not be used as a taxonomically relevant character in catenulids.

### New insights into catenulid phylogeny

In our previous study, *Stenostomum* was split into four clades (Tratkiewicz et al. 2026), for which distinct morphological traits can be identified that reflect phylogenetic position. The newly sequenced species obtained in the course of this study aligned with the previously characterized clades. For instance, *S. corderoi* exhibits all the characteristics of Clade 3: the absence of refracting bodies, a V-shaped mouth with lips, and closed sensory pits. Furthermore, *S. arevaloi* (previously an isolated lineage) now groups with *S. paraguayense* and *S. glandulosum* in Clade 5. This clade also shares common features, such as multiple pairs of refracting bodies and open, slit-like sensory pits (the presence of these traits in *S. glandulosum* is documented by Kepner (1930), and in *S. arevaloi* by (Marcus 1945)). Consequently, the systematics of the genus *Stenostomum* have largely stabilised, with five distinguishable clades supported by both molecular and morphological data. The newly sequenced species fit neatly into these clades, aligning with their respective morphological characteristics.

Another notable result of our study concerns the phylogenetic relationships within the family Catenulidae. We recorded animals which we identified as *Suomina evelinae*. Due to the presence of a statocyst, a ciliated ring around the mouth, and characteristic round epidermal inclusions, we are confident in our identification of these specimens as *S. evelinae* sensu Marcus (1945). This species was first described by Marcus (1945) and placed in the genus *Suomina* alongside *S. turgida* and *S. sawayai*. Later, Larsson and Jondelius (2008) moved *Suomina* into *Catenula*, based on the molecular data from *C. turgida*, which was recovered in a monophyletic position within the *Catenula* clade. In the present study, *C. turgida* was retrieved in a similar position within *Catenula*; however, *S. evelinae* grouped closer to *Paracatenula*. Based on the original description of *Suomina* and the retrieved phylogenetic position of this species, we propose restoring the combination *Suomina evelinae*. Elucidating the exact relationships within the family Catenulidae will, however, require denser sampling of the other genera on this branch of the catenulid tree – particularly the sequencing of genera currently lacking any molecular data, such as *Dasychormus*, *Chordarium* and *Africatenula*, as well as additional species of *Catenula* and *Suomina*.

We also speculate on the similarity between our specimens of *S. evelinae* and *Dasychormus pygmaeus* recorded by Kolasa (1973). The original description of this species was provided by Reisinger (1924) as *Catenula pygmaeus*. Although the description is vague and lacks drawings, Reisinger reports characteristics that we also observed in our specimens, including a statocyst, a ring of cilia, a wrinkled tail, and asexual reproduction with a maximum of two zooids. Marcus (1945) later described *Dasyhormus lithophorus* and noted its similarity to *C. pygmaeus*, though the absence of drawings and the different habitat of those specimens make the comparison uncertain. The third and final description of what is now *Dasyhormus pygmaeus* was provided by Kolasa (1973), in his botanic garden studies. The similarity between our specimens and Kolasa’s drawing, combined with the comparable habitat (*Sphagnum* samples from an exotic greenhouse in both cases), leads us to question the correct identification of this *Dasyhormus* record. Given the vague original description of *D. pygmaeus* and its sparse records — particularly the lack of reliable drawings or photographic documentation — we consider this species a nomen dubium. Nevertheless, recording and sequencing of *D. lithophorus* as described by Marcus could help determine its relationships within Catenulida and clarify the exact phylogenetic relationship between *S. evelinae* and the genus *Dasyhormus*.

## Conclusions

Greenhouses, regarded as tropical islands in temperate zones, can serve as a source of non-native animal species characteristic of warmer habitats, and our results confirm that they represent a source of exotic species for barcoding and taxonomic studies. One limitation of this type of research is establishing the origin of species, which can be especially difficult for taxa with limited biogeographic data, such as microturbellaria. Nevertheless, specimens obtained from botanic gardens can provide valuable material for taxonomic work on microturbellaria, as demonstrated here. Overall, our data suggest that in temperate climates, sampling in greenhouses is a simple and cost-effective complement to traditional field studies, particularly for barcoding of microscopic invertebrates.

## Additional information

### Ethical statement

No ethical statement was reported.

### Artificial Intelligence (AI) use

The authors accept full responsibility for the content of the manuscript, including the disclosure of any use of AI.

No AI tools were used in the preparation of this manuscript.

### Author contributions

K.T. collected, photographed, and identified the animals, cultured the animals, performed molecular laboratory work, processed the sequences, performed the phylogenetic analyses, prepared figures, and drafted the manuscript. M.S. collected, photographed, and cultured the animals. M.Z. provided the study material. L.G. designed the study, acquired funding, photographed and identified the animals, and reviewed and edited the manuscript. All authors read and accepted the final version of the manuscript.

### Data availability

All of the data that support the findings of this study are available in the main text or Supplementary Information. The molecular data underlying this article are available in GenBank under accession numbers PZ143194-PZ143204; PZ164137-PZ164143 and PZ210113-PZ210119.

